# SARS-CoV-2 ORF8 can fold into human factor 1 catalytic domain binding site on complement C3b: Predict functional mimicry

**DOI:** 10.1101/2020.06.08.107011

**Authors:** Jasdeep Singh, Sudeshna Kar, Seyed Ehtesham Hasnain, Surajit Ganguly

## Abstract

Pathogens are often known to use host factor mimicry to take evolutionary advantage. As the function of the non-structural ORF8 protein of SARS-CoV-2 in the context of host-pathogen relationship is still obscure, we investigated its role in host factor mimicry using computational protein modelling techniques. Modest sequence similarity of ORF8 of SARS-CoV-2 with the substrate binding site within the C-terminus serine-protease catalytic domain of human complement factor 1 (F1; PDB ID: 2XRC), prompted us to verify their resemblance at the structural level. The modelled ORF8 protein was found to superimpose on the F1 fragment. Further, protein-protein interaction simulation confirmed ORF8 binding to C3b, an endogenous substrate of F1, via F1-interacting region on C3b. Docking results suggest ORF8 to occupy the binding groove adjacent to the conserved “arginine-serine” (RS) F1-mediated cleavage sites on C3b. Comparative H-bond interaction dynamics indicated ORF8/C3b binding to be of higher affinity than the F1/C3b interaction. Hence, ORF8 is predicted to inhibit C3b proteolysis by competing with F1 for C3b binding using molecular mimicry with a possibility of triggering unregulated complement activation. This could offer a mechanistic premise for the unrestrained complement activation observed in large number of SARS-CoV-2 infected patients.

## 1. Introduction

The world is currently facing an unprecedented pandemic caused by the transmission of a novel corona virus, named as severe acute respiratory syndrome coronavirus 2 or SARS-CoV-2 [1, 2]. Following the identification of the virus, the early phase of clinical characterization of the disease revealed that a substantial number of the patients progresses towards acute respiratory distress syndrome (ARDS) eventually leading to multi-organ failure [3-6]. Though the sequence of events leading to multiple organ failure is still not ascertained, it appears that the virus infection-induced hyperactivation of the complement system along with robust pro-inflammatory responses, vascular thrombus formation and coagulation could play a major role [7-10].

An emerging hypothesis suggests that the SARS-CoV-2 non-structural ORF8 (Open reading frame 8) protein could participate in immune evasion strategy [11]. However, it does not explain the tissue damaging immune storm that accompanies the pathogenesis. This led us to investigate whether the structural elements of ORF8 can mimic any human host factor that can elicit robust proinflammatory responses as a bystander effect. Diverse pathogens, including viruses, are believed to use molecular mimicries that resemble host factors to acquire evolutionary advantage [12, 13]. Towards this goal, we observed that ORF8 has a modest level of amino acid sequence similarity with the substrate binding site located within the C-terminal domain of human complement factor 1 (F1; PDB ID: 2XRC; Ref. 14). Here, we report that ORF8 has optimal structural resemblance (tertiary fold architecture) that might allow it to interact with one of the F1 substrates - human complement C3b, by adopting F1 mimicry. Using protein-protein interaction modelling, we predict ORF8 to bind to C3b via a site that appears to overlap with the F1 interacting interface on C3b. The possibility of competitive sequestration of F1 from C3b binding interface could form the basis of uncontrolled complement activation in the host.

## 2. Results

### 2.1. Modelling of SARS-CoV-2 ORF8 protein

Searching (Blastp suite, NCBI) the human (Taxid 9606) PDB protein database using the full length 121 amino acid long SARS-CoV-2 ORF8 protein did not generate any significant hits with E (expected threshold) value set at 1.0. However, increasing the E to 10 (default) revealed a lower level of sequence similarity of ORF8 (48% similarity, 25% identity, E value 1.4) with C-terminal domain (amino acid residues 500 to 557) of F1. The resemblance of the ORF8 sequence was with the substrate binding region of F1, embedded within the catalytic domain, as schematically shown in Figure 1A. According to the NCBI Conserved Domain database, F1 protein is a trypsin-like serine protease with the domains as described (Figure 1A). The catalytic domain extends from 322 to 557 residues after the zymogen cleavage site between R321 and I 322 residues. This F1 catalytic domain is subdivided into an active site, composed of consensus triad residues - H 363, D 411, and S 507; and, a substrate binding site from residues D 501 to R 557 [15]. In comparison, these triad catalytic residues are missing in ORF8. However, it has resemblance from E 57 (corresponding D 501 in F1) to R 115 (corresponding R 557 in F1) in the substrate binding region (Figure 1A).

**Figure 1.**
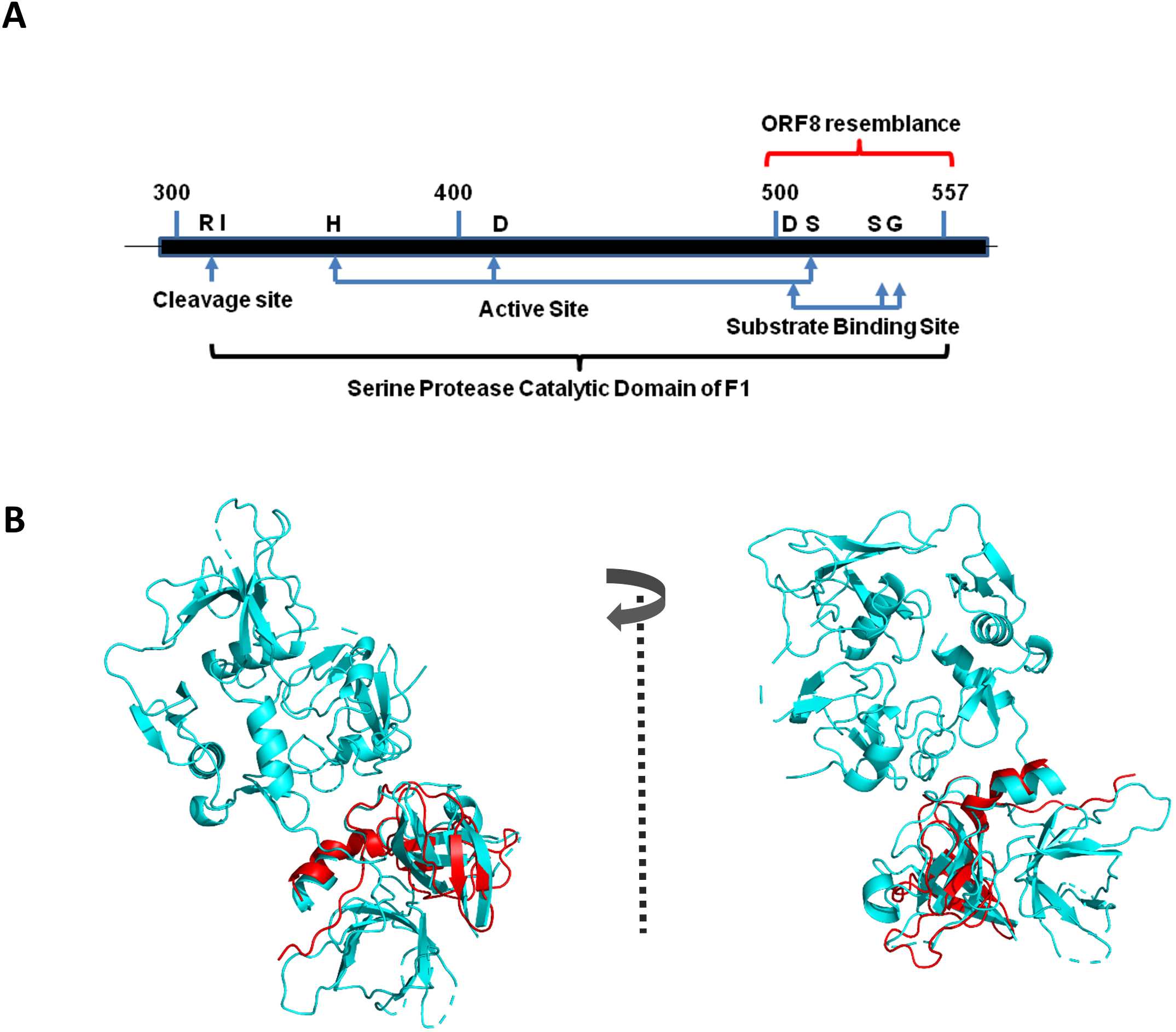
**A**. Cartoon showing the region of F1 (pdb 2XRC_A / gi 339961198) that has sequence resemblance with ORF8. **B**. Structural superimposition of modelled ORF8 (residues 18-112) of SARS-CoV-2 (Red) with C-terminal domain (residues 421-554) of F1 (PDB id: 2xrc) in Cyan as template. Clockwise rotation provides frontal and rear view of the overlapping residues.

Modelling of ORF8, followed by overlaying it on the template F1 (PDB id: 2xrc), the sequence similarity was reflected at the structural level as well (Figure 1B). The α-helix at the C-terminus portion of ORF8 (Red ribbon) linked via a central large loop to the β-sheet superimposes with the helix-loop-β-sheet portions of the C-terminus catalytic domain of F1 (Cyan ribbon). Taken together, the sequence resemblance of ORF8 is also reinforced at the structural level as it could adopt tertiary fold architecture of F1 catalytic domain.

### 2.2. Interaction modelling of ORF8 with Complement C3B

Human complement C3b has been regarded as one of the major substrates of F1 [14, 16]. F1-mediated cleavage of C3b is believed to turn-off complement activation and act as a negative-feedback regulatory step. Since ORF8 has a structural similarity with the substrate binding portion enclosed within the C-terminal catalytic domain of F1, we determined the ability of ORF8 of SARS-CoV-2 to compete with F1 protein for its binding with C3b (PDB Entry - 2I07). To accomplish this, we applied a protein-protein docking algorithm (unbiased rigid docking solution) where no pre-defined interaction sites were assigned. The top ranked solutions from both docked complexes showed partial overlap of C3b binding site for ORF8 and F1 occupancy (Figure 2A and 2B). Interestingly, the global energy for top ranked solution of C3b-ORF8 docking (−23.16) was slightly higher than C3b-F1 docking (−20.85). ORF8 interactions with C3b were found to occur via multiple hydrogen bonding with residues including N 939, H 1349, Y 1482 and others (Figure 2B; Tables 1 and 2). These interacting residues are encompassed by F1 interacting region (Figure 2A) on chain B of C3b. Furthermore, ORF8 appears to be docked in close proximity to the R 1303 / S1304 and R1321 / S 1322 (Blue spheres in Figure 2B), the target of F1 catalytic site for proteolysis. Moreover, the lack of conserved serine protease catalytic triad residues H, D and S, as present in F1, might allow ORF8 to act like a competitive inhibitory ligand, hindering access to F1 binding. Thus, ORF8 might subsequently shield proteolytic cleavage between residues R 1303 and S 1321 (Figure 2A) or other sequential cleavage sites on C3b CUB (C1r/C1s, Uegf, Bmp1) domains, as described previously [14], with a potential to hinder C3b down-regulation and attenuate opsonin generation.

**Table 1:** Protein protein interaction network between pre-simulated F1 and C3b. Residues involved in inter-molecular H-bonds and salt-bridge formation are indicated.

| H-bonds |  |
| --- | --- |
| C:ARG 430 [NH2] | B:GLU 769 [OE1] |
| C:SER 256 [O] | B:TYR1428 [HH ] |
| C:GLU 270 [O] | B:TYR1447 [HH ] |
| C:GLN 373 [OE1] | B:GLN 983 [NE2] |
| C:THR 377 [O] | B:ARG 979 [NH2] |
| C:ARG 388 [O] | B:ARG 979 [NH2] |
| C:ASP 447 [OD2] | B:TYR1482 [HH] |
| C:THR 448 [OG1] | B:TYR1483 [HH] |
| Salt bridges |  |
| C:ARG 430 [NH2] | B:GLU 769 [OE1] |
| C:GLU 470 [OE1] | B:LYS1360 [NZ] |

**Table 2.** Protein-protein interaction network between ORF8 and C3b. Residues involved in inter-molecular H-bonds and salt-bridge formation are indicated.

| H-Bonds |  |
| --- | --- |
| A:ILE 58[ N ] | B:TYR1482[ OH ] |
| A:ASP 107[ N ] | B:GLU1487[ OE2] |
| A:SER 21[ O ] | B:HIS1349[ NE2] |
| A:SER 24[ OG ] | B:ASN 939[ ND2] |
| A:GLU 106[ OE1] | B:ASN1484[ ND2] |
| A:ASP 107[ OD2] | B:LYS1360[ N ] |
| A:ASP 113[ O ] | B:GLU1486[ N ] |
| Salt-bridges |  |
| A:LYS 2[ NZ ] | B:ASP 832[ OD2] |
| A:HIS 28[ NE2] | B:GLU1486[ OE1] |
| A:ASP 113[ OD1] | B:LYS1478[ NZ ] |
| A:ASP 113[ OD2] | B:LYS1478[ NZ ] |

**Figure 2.**
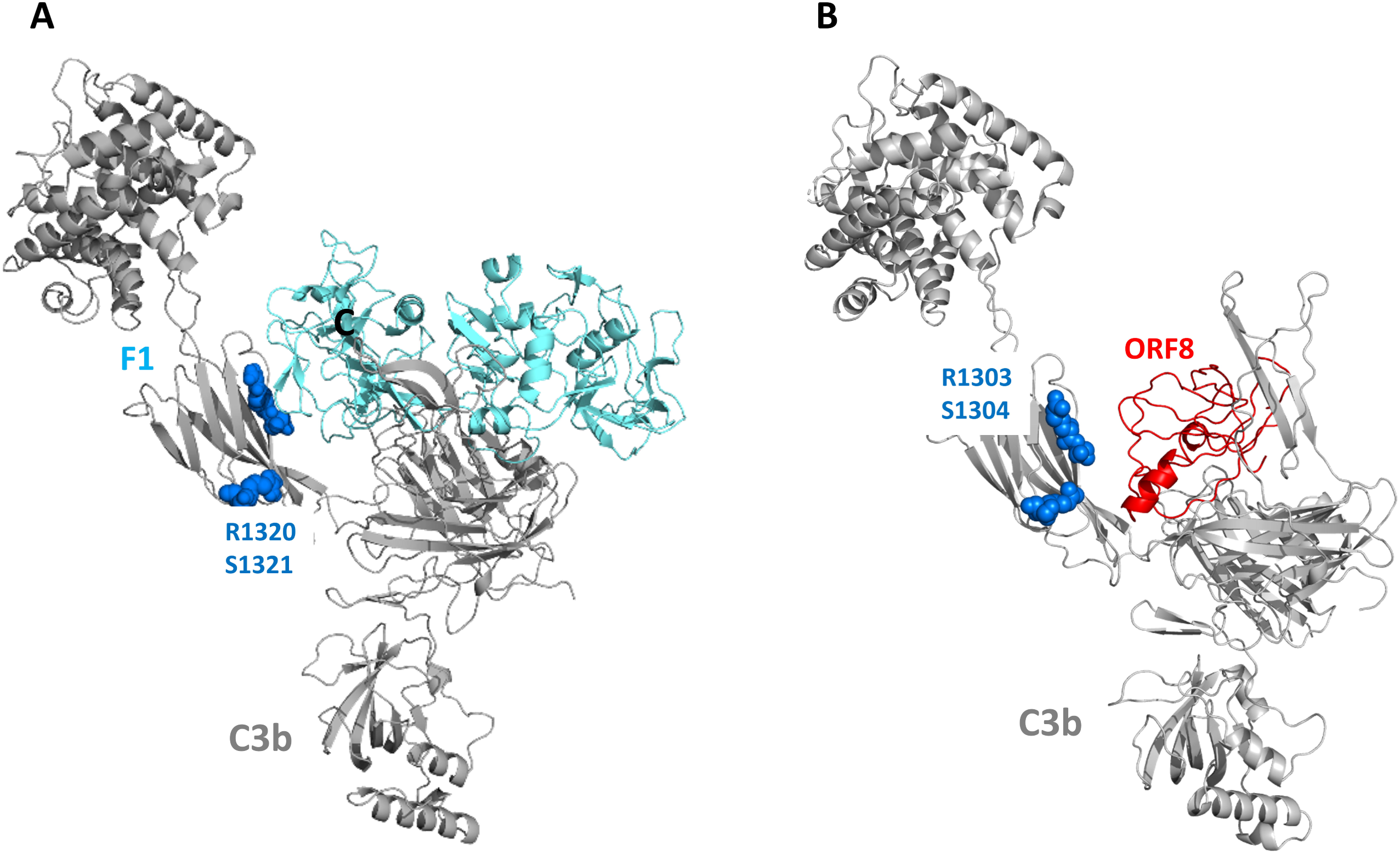
Modelling and Protein-protein docking of ORF8/F1 complement with C3b human complement. A. Highest ranked model obtained from protein-protein docking of F1 complement (Cyan) and human C3b complement (Grey). B. Highest ranked model obtained from protein-protein docking of ORF8 (Red) and human C3b complement (Grey). Blue spheres indicate Arg-Ser (RS) protease cleavage sites encompassing C3b CUB (C1r/C1s, Uegf, Bmp1) domain near the docked complexes. The arrows from (A) define outcomes from protein-protein docking of C3b with human F1complement (Cyan arrow) and ORF8 (Red arrow).

### 2.3. Validation of ORF8 and C3b interaction accuracy

For validation of the accuracy of the docking method adopted and in an effort to minimize the false positive docking solutions, we employed two approaches. In the first approach, we used a mutant form of ORF8, mORF8 bearing amino acid substitutions S24L, V62L and L84S. These mutations were found naturally in viral isolates (Supplementary Fig. S1). Mutation induced structural stability analysis displayed S24L and V62L substitutions to have minor stabilization effects on ORF8 architecture (ΔΔG_wild-type→mORF8_ +0.36 and +0.09 kcalmol^-1^, respectively). In contrast, the high entropy (globally dominant) L84S mutation showed much higher destabilization effect on ORF8 (ΔΔG_wild-type→mORF8_ −1.07 kcalmol^-1^) (Supplementary Fig.2). In the top ranked solution for mORF8-C3b docking, mORF8 was bound to a site different from F1 and wild-type ORF8 interacting interfaces (Global energy - 15.82) (Figure 3A and 3B). However, the docking solution which showed binding of mORF8 at the proximity of F1 and ORF8 binding site yielded a much lesser global energy score (−2.9) (Figure 3A and 3C). This suggests that by rigidly fitting the mORF8 in the F1-C3b interface would make the binding energetically less favourable. Although both ORF8 and mORF8 were structurally conserved (RMSD ∼0.2 Å; Figure 3A), the mutations have clearly perturbed mORF8 binding with C3b. It appears that the preferential interaction interface of mORF8 with C3b are dissimilar from F1-C3b binding interface, indicating that mORF8 is less likely to compete with F1 for the same binding site on C3b. Thus, it appears that the wild type ORF8 conformation is required for precisely fitting into the binding groove of F1-C3b interface.

**Figure 3.**
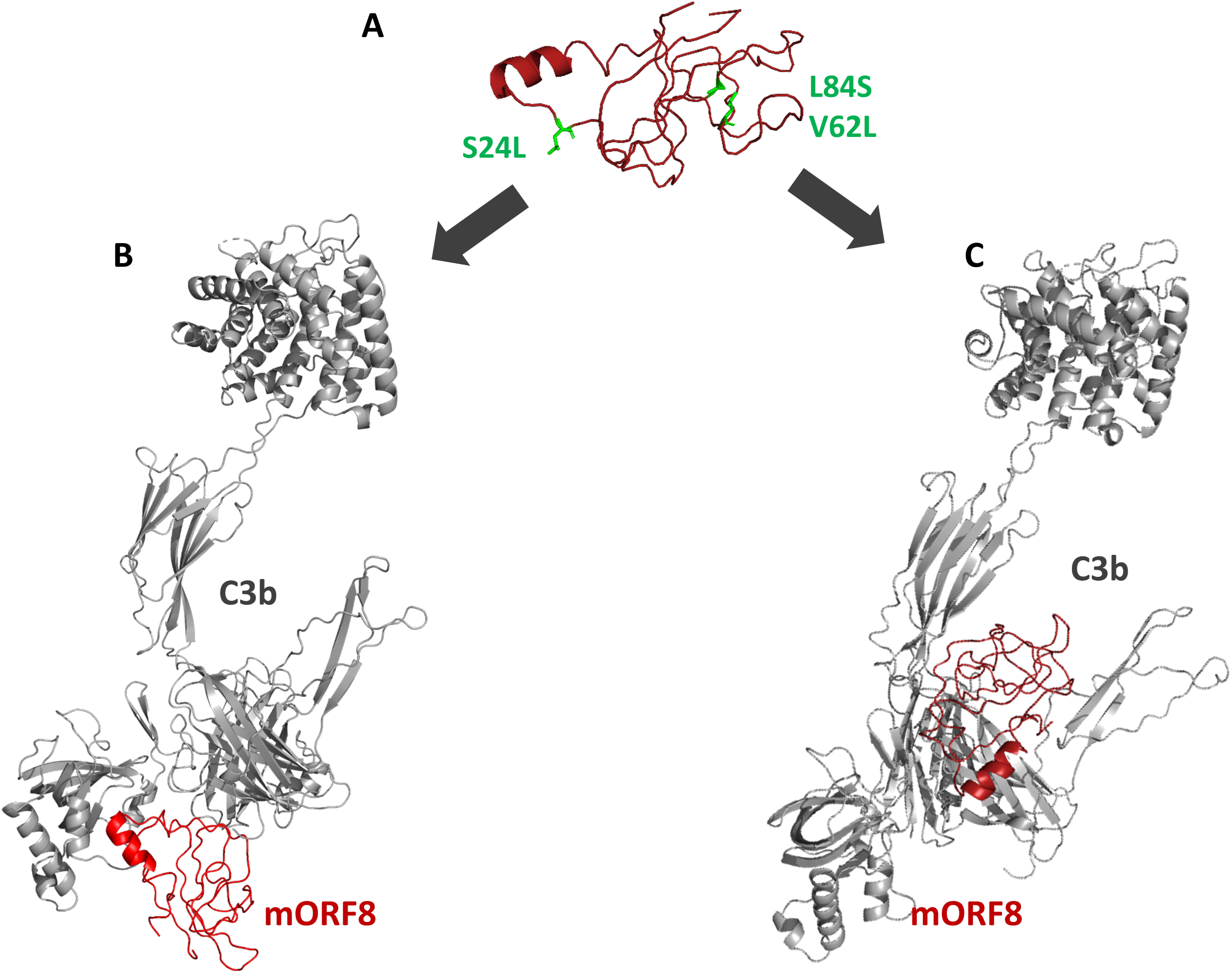
Modelling and Protein-protein docking of mORF8 with C3b human complement. **A**. Modelled ORF8 (Brown) showing three high entropy mutations; S24L, V62L and L84S (Green Sticks). **B**. Highest ranked model obtained from protein-protein docking of mORF8 (Brown) and human C3b complement (Grey). **C**. Low energy model obtained from protein-protein docking of mORF8 (Brown) and human C3b complement (Grey) at F1 binding site. The arrows (Black) define outcomes from protein-protein docking of C3b with mORF8 at its two different sites.

In the second validation approach, we used two randomly selected SARS-CoV-2 proteins – ORF7a (121 residue length; green) protein and 1 - 121 amino acid fragment of NSP10 (orange) of SARS-CoV-2, with matching amino acid length as ORF8, as negative controls for modelling interactions with C3b (Figure 4). The top ranked docking results for both proteins show surprising associations with C3b at a common region between amino acid residues 771 – 850. The interacting regions appear to be located at an exterior surface, away from the ORF8/F1 binding groove on the C3b protein. Anti-parallel β–sheets sometimes provide stable binding interface for protein-protein interactions [17]. Since both unrelated proteins bind to same site on C3b supported by two partial anti-parallel sheets, it appears that these docking solutions are more of a reflection of algorithmic limitations than representing physiological states. In contrast, the binding of ORF8 with C3b is driven by structural resemblance with F1, and is a testimony to protein mimicry action. It is to be noted that in spite of the projected higher affinity of ORF8 for a shared docking region on C3b, the effort here is to identify the “near-native” conformation that dictates the physiological impact of the ORF8-C3b complex. The top-ranked solution does not always replicate the physiological states. Hence, in the absence of crystal structure of the native ORF8, these docking solutions only present a comparative evaluation of ORF8 in the context of F1 mimicry functions.

**Figure 4.**
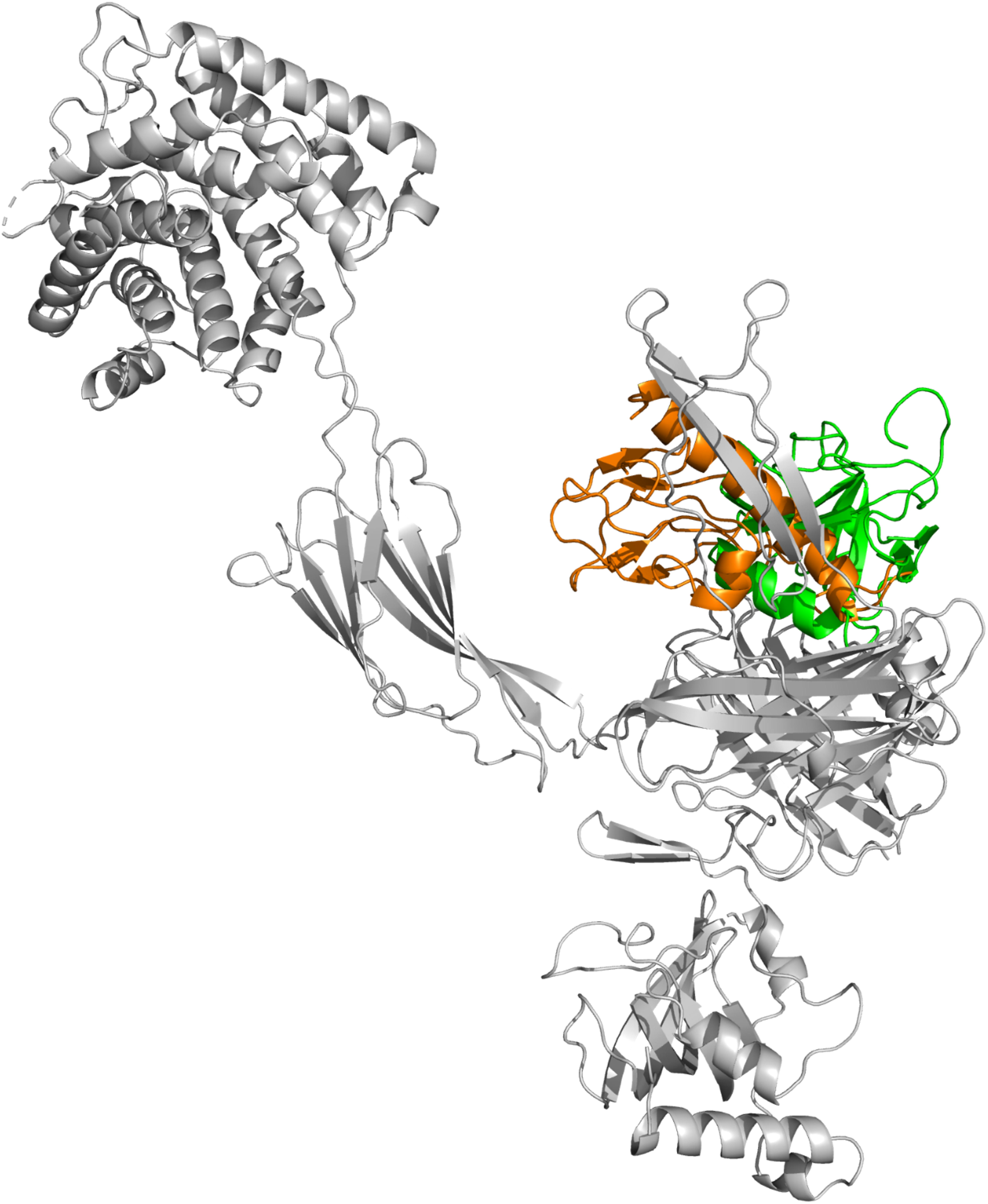
Protein-protein docking of C3b (Grey) with a structurally unrelated 121 amino acid ORF7a (Green) and NSP10 (Orange). Both proteins showed interaction sites between residues 771-850, different from F1 and ORF8 interactions with C3b.

### 2.4. Comparative dynamics of ORF8-C3b versus F1-C3b complex

To understand the dynamics of F1/ORF8 interactions with C3b, we performed MD simulations of both complexes for 25 ns. Variation in H-bond interaction network for both systems showed higher number of H-bonds/polar contacts between ORF8-C3b (21.09 H-bonds per time frame) compared to F1-C3b (18.45 H-bonds per time frame) (Figure 5). This further reinforced our docking outcomes of comparatively higher affinity binding of ORF8 to the same site where F1 binds. Thus, it appears that ORF8 possesses the capability of competing with F1 for a common binding pocket on C3b.

**Figure 5.**
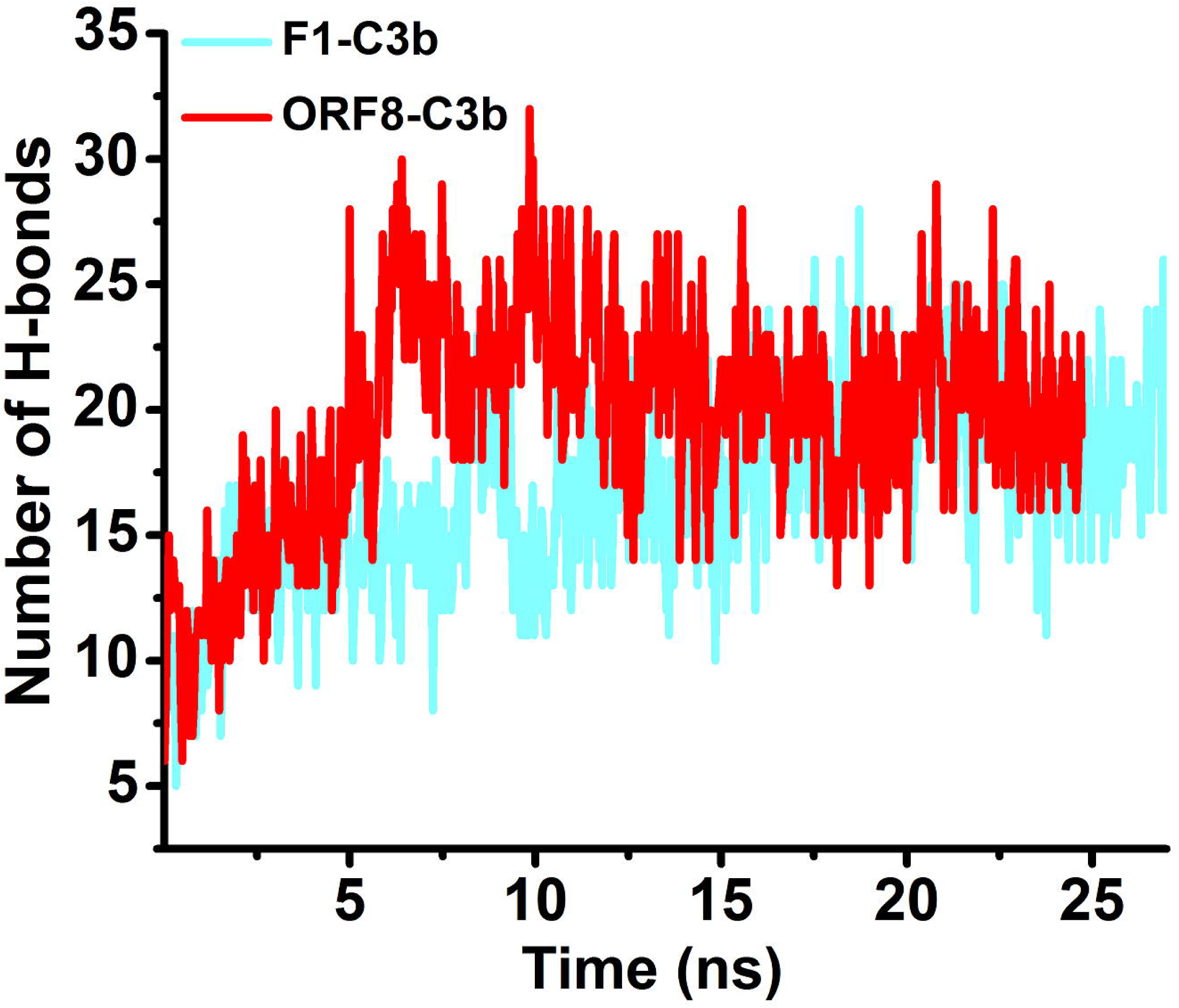
MD simulations of top ranked F1-C3b and ORF8-C3b docked complexes. Evolution of H-bonds between F1 factor and C3b complement complex (Cyan) and ORF8-C3b complex (Red) over the simulation period of 25 ns.

## 3. Discussion

The current study highlights immunomodulatory potential of ORF8 protein of SARS-CoV-2 through molecular mimicry of a portion of F1 protein, the negative-feedback regulator of complement pathway. Using a combination of MD simulations and protein-protein docking approach, we have compared binding interactions of ORF8 and F1 proteins with C3b. Our results predict a plausible mechanism of infection-induced host complement hyperactivation coupled with evasion strategy adopted by SARS-CoV-2 through sustained protection of C3b protein via competitive disruption of F1 binding.

F1 is known to function as a negative regulator of the host complement system by regulating the levels of C3b complexes [14, 16, 18]. In this context, the resemblance of SARS-COV-2 ORF8 protein with the substrate binding domain of human F1 is intriguing. It is possible that ORF8 protein can directly interact with active complement factors like C3b mimicking F1 binding. As ORF8 possesses only similar substrate binding motif and lacks the catalytic triad residues of F1, this perhaps allows ORF8 to act like a competitive inhibitor peptide, rapidly blocking F1 access to the target proteolytic sites on C3b. The prevention of C3b proteolysis has two major consequences in the host – (a) uncontrolled accumulation of C3b which might lead to sustained complement stimulation (b) absence of C3 proteolysis blocks the production of opsonins products like ic3b, c3dg and c3d, helping the virus to escape immune surveillance. Both fates are detrimental to the host.

Complement cascade activation in response to viral infections can have a pathologic as well as protective effect in the host. For example, C3 activation is fundamental in viral clearance and host protection in H1N1 and Asian Avian H5N1 influenza A viruses [19]. In contrast, C3 forms the foundation of SARS-CoV–induced disease pathogenesis as evinced by C3^-/-^ mice exhibiting significantly attenuated virus-induced systemic inflammation and tissue injury [20]. In cases where the host complement system colludes with the viral pathogens to exacerbate disease pathogenesis, amplification of the C3b-C3 convertase reaction loop, leading to the formation of master inflammatory mediator C5a and tissue-damaging C5b, overpowers the C3b breakdown cycle by its negative regulators. F1 is one of the main proteases that degrade C3b, thus acting as a molecular brake to maintain and regulate complement homeostasis. Disruption of F1-mediated inactivation of C3b would lead to uncontrolled complement cascade stimulation, leading to aggravated chronic inflammation, tissue damage and ultimately, multiorgan-failure and death. Since an overactive and maladapted host complement system is shown to be a requisite component of SARS-CoV-2 infection and a determining factor in the severity and progression of the illness [7], we can hypothesize that the molecular mimicry function of ORF8 serves to shield C3b from FI-mediated proteolytic degradation and perpetuate a feed-forward loop of C3b-C3 convertase interaction. At the same time, ORF8 binding to C3b and shielding the F1 cleavage sites in the CUB domain of C3b bears the possibility of limiting the generation of C3b proteolytic products, which act as opsonins and could be detrimental to virus survival. Thus, if C3b is shielded from degradation, it might allow the SARS-CoV-2 to escape the complement surveillance.

## 4. Conclusion

Recognition and elimination of viral particles by the host complement system has been known since 1930 [21]. During the adoption of humans as host, viruses have also evolved unique strategies to subvert the complement mediated innate immunity [22-24]. Encoding orthologs of the complement family of proteins or co-opting the host complement system are documented extensively [25]. Our results predicting the use of molecular mimicry as a strategy by SARS-CoV-2 to deregulate complement system is one such action designed to gain advantage during infection. The prediction of ORF8 as a binding partner of the C3b and competitively blocking the actions of F1, the negative regulator of complement homeostasis, could serve as an intriguing hypothesis explaining the immune surge in SARS-CoV-2 infected patients. Thus, in spite of the absence of experimental correlations, the current work presents stimulating insights into functional role of ORF8 at host-viral interface and warrants further investigation.

## 5. Materials and Methods

### 5.1. NCBI Reference Sequences

The 121 amino acid residues long ORF8 protein, NCBI reference sequence YP_009724396.1, was used for the entire study. This protein sequence was called wild type. For analysis with mutant protein (mORF8), triple substitutions S24L, V62L and L84S were incorporated in the wild type ORF8 sequence representing natural mutations as demonstrated in Supplementary figure S1, was used. Randomly selected SARS-CoV-2 derived full-length 121 amino acid long ORF 7a (YP_009724395.1) and 1-121 resides (c-terminal truncated) of NSP10 (YP_009725306.1) with an intention to match ORF8 amino acid length, were used as negative controls.

### 5.2. Methods for protein modelling and protein-protein docking

Modelling of wild type (NCBI reference genome) ORF8 and mutant ORF8 (mORF8; triple mutations S24L, V62L and L84S as described in (Supplementary fig. S1) was performed using I-TASSER (Iterative Threading ASSEmbly Refinement) with human complement factor 1 (F1; PDB id: 2xrc) as template. The stereo chemical quality of resulting model was verified using Ramachandran and ProSA analysis. Without assigning any prior binding site (unbiased), protein-protein dockings were performed employing Patchdock using clustering RMSD of 4.0 and subsequently top 20 resulting solutions were refined using Firedock (26 - 30). The global energy of top 20 ranked solutions by FireDock were calculated from contribution of attractive and repulsive vander waals forces, atomic contact energy (ACE) and contribution of the hydrogen bonds. Interaction analysis was carried out using PDBPisa (https://www.ebi.ac.uk/pdbe/pisa/). Effects of mutations on the structural stability of ORF8 was predicted using Dynamut (http://biosig.unimelb.edu.au/dynamut/).

### 5.3. Molecular dynamics simulations and setup

MD simulations were carried using gromacs v2016. Simulations of F1-C3b and ORF8-C3b complexes were carried at 300 K in a cubical box with 10 nm spacing from box edges. Ionic concentrations was kept at 0.1 M NaCl followed by system minimization by Steepest descent protocol. Equilibration and production runs (25 ns) were carried using parameters detailed in our previous reports [31, 32]. Images were constructed using PyMol while data was analysed using standard gromacs tools.

## Acknowledgement

The authors acknowledge the sacrifices of all those who are fighting the Covid-19 pandemic. JS acknowledges financial support under Young Scientist scheme of Dept. of Health Research (DHR), Ministry of Health and Family Welfare, Government of India. SEH is a JC Bose National Fellow of the Department of Science & Technology (DST), Government of India. SEH is a Robert Koch Fellow of the Robert Koch Institute. Berlin, Germany. SK acknowledges SERB/DST for core research support. SG thanks DHR, ICMR and SERB/DST for research support.

