## Supplemental Figs S1 and S2 for "SARS-CoV-2 ORF8 can fold into human factor 1 catalytic domain binding site on complement C3b: Predict functional mimicry"

**Supplementary Figures**

**Figure S1:** Shannon entropy as a measure of variation protein sequence alignment of 1042 sequences of ORF8. The amino acid variation plot is used to calculate the entropy at each position in a sequence set. Amino acids at position 24 (S24L), 62 (V62L) and 84 (L84S) show significant divergence and mutations as mentioned.

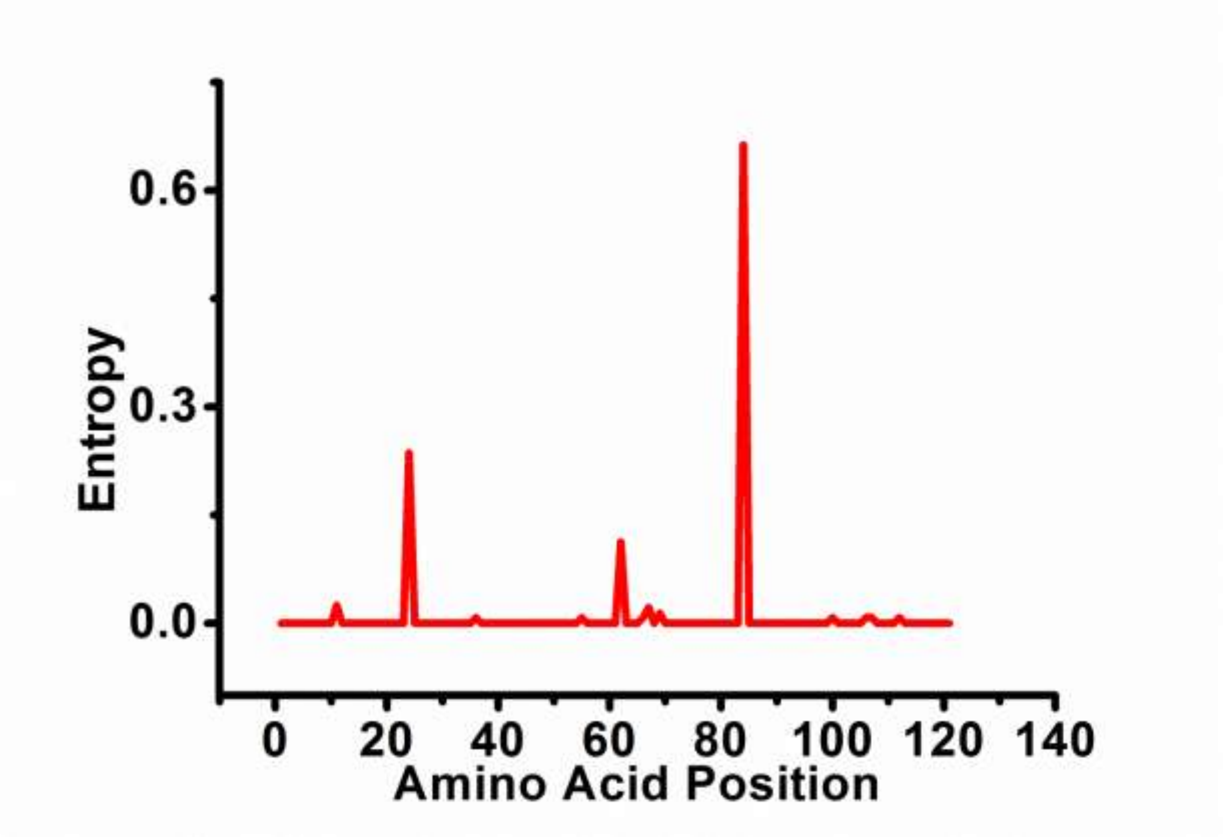

**Figure S2.** Residue interaction network of (A) Leu84 (wild type) and (B) Ser84 (Mutant). L84S mutation in ORF8 protein leads to decrease in strong H-bond and weak H-bond network around mutated residue.

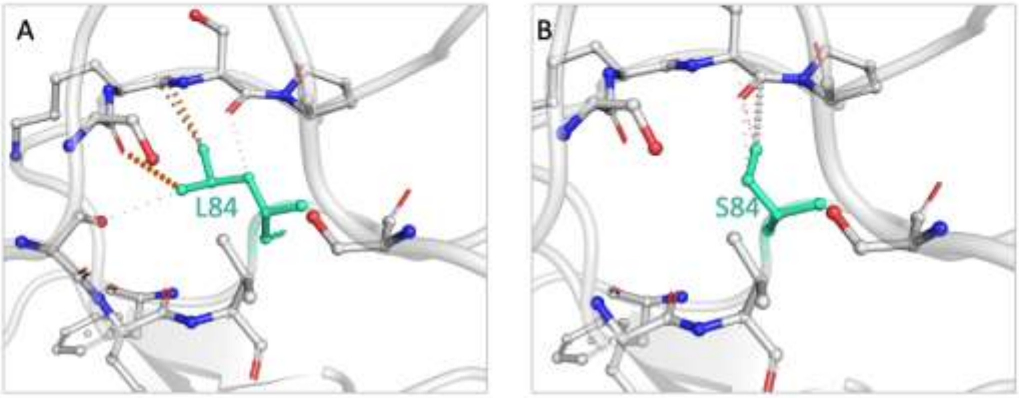
